# OPUS-CSF: A C-atom-based Scoring Function for Ranking Protein Structural Models

**DOI:** 10.1101/163972

**Authors:** Gang Xu, Tianqi Ma, Tianwu Zang, Qinghua Wang, Jianpeng Ma

## Abstract

We report a <u>C</u>-atom-based <u>s</u>coring <u>f</u>unction, named OPUS-CSF, for ranking protein structural models. Rather than using traditional Boltzmann formula, we built a scoring function (CSF score) based on the native distributions (analyzed through entire PDB) of coordinate components of mainchain C atoms on selected residues of peptide segments of 5, 7, 9, and 11 residues in length. In testing OPUS-CSF on decoy recognition, it maximally recognized 257 native structures out of 278 targets in 11 commonly used decoy sets, significantly more than other popular all-atom empirical potentials. The average correlation coefficient with TM-score was also comparable with those of other potentials. OPUS-CSF is a highly coarse-grained scoring function, which only requires input of partial mainchain information, and very fast. Thus it is suitable for applications at early stage of structural building.

## Introduction

A potential function plays a central role in predicting protein structures. Generally, there are two kinds of potential functions: physics-based potentials and knowledge-based potentials. Physics-based potentials typically are the all-atom molecular mechanics force-fields (Arnautova et al., 2006; Brooks et al., 1983; Case et al., 2005; MacKerell Jr et al., 1998; Weiner et al., 1986), such as CHARMM (Brooks et al., 1983; MacKerell Jr et al., 1998) and AMBER (Case et al., 2005). They also include coarse-grained potentials such as MARTINI (Marrink et al., 2007), UNRES (Liwo et al., 1997a; Liwo et al., 1997b) and OPEP (Chebaro et al., 2012).

The knowledge-based potentials are derived from statistical analysis of known structures and are widely used in structural prediction (Buchete et al., 2004; DeBolt and Skolnick, 1996; Gilis et al., 2006; Gohlke and Klebe, 2001; Hendlich et al., 1990; Hoppe and Schomburg, 2005; Jernigan and Bahar, 1996; Jones et al., 1992; Koliński and Bujnicki, 2005; Lazaridis and Karplus, 2000; Lu and Skolnick, 2001; Lu et al., 2008; Ma, 2009; Miyazawa and Jernigan, 1985; Moult, 1997; Poole and Ranganathan, 2006; Russ and Ranganathan, 2002; Samudrala and Moult, 1998; Shen and Sali, 2006; Sippl, 1990, 1995; Skolnick, 2006; Skolnick et al., 2000; Tobi and Elber, 2000; Wu et al., 2007; Yang and Zhou, 2008; Zhang et al., 1997; Zhang and Zhang, 2010; Zhang et al., 2003; Zhou and Skolnick, 2011; Zhou and Zhou, 2002; Zhou et al., 2006). In general, knowledge-based potentials can be either constructed at coarse-grained residue level (Buchete et al., 2004; Gilis et al., 2006; Hendlich et al., 1990; Hoppe and Schomburg, 2005; Jones et al., 1992; Koliński and Bujnicki, 2005; Miyazawa and Jernigan, 1985; Sippl, 1990; Skolnick et al., 2000; Tobi and Elber, 2000; Wu et al., 2007; Zhang et al., 2003) or at atomic level (DeBolt and Skolnick, 1996; Lu and Skolnick, 2001; Lu et al., 2008; Samudrala and Moult, 1998; Shen and Sali, 2006; Yang and Zhou, 2008; Zhang et al., 1997; Zhang and Zhang, 2010; Zhou and Skolnick, 2011; Zhou and Zhou, 2002). Although coarse-grained potentials may not be rigorous, it helps to focus on essential features and excludes less important details, thus reduces computational cost (Kmiecik et al., 2016; Noid, 2013). The performance of coarse-grained potential is highly related to how one designs the coarse-graining scheme. For example, OPUS-Ca potential (Wu et al., 2007) uses the positions of Ca atoms as input, calculate other atomic positions as pseudo-positions and significantly reduces the computing cost. Other applications of coarse-grained models using Ca positions are also reported in literature (Wu et al., 2005a; Wu et al., 2005b).

In this work, unlike traditional empirical potential functions using Boltzmann formula, we built a scoring function based on the native distributions of coordinate components of mainchain C atoms on a few selected residues of small peptide segments of 5, 7, 9 and 11 residues in length. A lookup table was first generated for native distributions of coordinate components by analyzing peptide segments in the entire Protein Data Bank (PDB). Then the scoring function was calculated for a particular test structure by comparing the information of its segments with the lookup table. The performance of OPUS-CSF was tested on 11 commonly used decoy sets, the results indicated that OPUS-CSF recognized significantly more native structures from their decoys than other empirical potentials. In terms of the correlation coefficients between CSF scores and TM-scores, they were comparable with those of the popular all-atom empirical potentials. Most importantly, OPUS-CSF achieved such performance despite its highly coarse-grained nature. That indicates the advantages of OPUS-CSF in terms of its speed and its application in the early stage of structural modeling. This is vitally important for applications such as building structural models against intermediate resolution data from experimental techniques like cryogenic electro-microscopy (cryo-EM).

## Results and Discussion

We compared the performance of OPUS-CSF on 11 commonly used decoy sets with that of popular all-atom potential functions. In Table 1, we listed the results of 5-residue segment case (OPUS-CSF5) and all-segment combined case (OPUS-CSF). For 5-residue segment case, OPUS-CSF5 successfully recognized 244 out of 278 native structures from their decoys and had the average Z-score (-3.56) comparable with that of GOAP (-3.57). For combined segment case, OPUS-CSF successfully recognized 257 out of 278 native structures from their decoys and had the average Z-score (-4.12) better than that of GOAP (-3.57). It is interesting that although OPUS-CSF is a highly coarse-grained scoring function, its performance is significantly better than other all-atom potentials.

**Table 1.** The results of OPUS-CSF5 (5-residue segment) and OPUS-CSF (combined segment length) on 11 decoys sets compared with different potentials. The results of other potentials come from GOAP paper. The numbers of targets, with their native structures successfully recognized by various potentials, are listed in the table. The numbers in parentheses are the average Z-scores of the native structures. The bigger the absolute value of Z-score, the better. Out of totally 278 targets in 11 decoy sets, OPUS- CSF5 (5-residue segment) recognized 244 and OPUS-CSF (combined segment length) recognizes 257 native structures from their decoys. The bold number in each row indicates the best one among all the potential functions for that particular decoy set (if the numbers of targets are the same, the bold face is on the one with the better Z-score).

| Decoy sets | Total # of targets | DFIRE | RWplus | dDFIRE | OPUS-PSP | GOAP | OPUS-CSF5 | OPUS-CSF |
| --- | --- | --- | --- | --- | --- | --- | --- | --- |
| 4state_reduced | 7 | 6(-3.48) | 6(3.51) | 7(-4.15) | **7(-4.49)** | 7(-4.38) | 7(-3.38) | 7(-3.31) |
| fisa | 4 | **3(-4.87)** | 3(-4.79) | 3(-3.80) | 3(-4.24) | 3(-3.97) | 2(-2.31) | 2(-2.55) |
| fisa_casp3 | 5 | 4(-4.80) | 4(-5.17) | 4(-4.83) | **5(-6.33)** | 5(-5.27) | 4(-4.38) | 4(-6.72) |
| hg_structal | 29 | 12(-1.97) | 12(-1.74) | 16(-1.33) | 18(1.87) | 22(-2.73) | **23(-2.07)** | 23(-2.06) |
| ig_structal | 61 | 0(0.92) | 0(1.11) | 26(-1.02) | 20(0.69) | 47(-1.62) | 49(-2.03) | **56(-2.14)** |
| ig_structal_hires | 20 | 0(0.17) | 0(0.32) | 16(-2.05) | 14(-0.77) | 18(-2.35) | 19(-2.19) | **20(-2.08)** |
| I-TASSER | 56 | 49(-4.02) | 56(-5.77) | 48(-5.03) | 55(-7.43) | 45(-5.36) | 55(-5.32) | **56(-6.39)** |
| lattice_ssfit | 8 | 8(-9.44) | 8(-8.85) | 8(-10.12) | 8(-6.75) | 8(-8.38) | 8(-9.56) | **8(-11.79)** |
| lmds | 10 | 7(-0.88) | 7(-1.03) | 6(-2.44) | 8(-5.63) | 7(-4.07) | 8(-5.47) | **8(-6.80)** |
| MOULDER | 20 | 19(-2.97) | 19(-2.84) | 18(-2.74) | 19(-4.84) | 19(-3.58) | **20(-3.18)** | 20(-3.16) |
| ROSETTA | 58 | 20(-1.82) | 20(-1.47) | 12(-0.83) | 39(-3.00) | 45(-3.70) | 49(-3.68) | **53(-4.53)** |
| Total | 278 | 128(-1.94) | 135(-2.13) | 164(-2.52) | 196(-2.86) | 226(-3.57) | 244(-3.56) | **257(-4.12)** |

We also calculated the Pearson’s correlation coefficients between CSF score andTM-score (Zhang and Skolnick, 2004) in all decoy sets. The results are shown in Table 2. OPUS-CSF has comparable average correlation coefficient with those of GOAP and OPUS-PSP despite the fact that OPUS-CSF is highly coarse-grained and the other two are all-atom potentials.

**Table 2.** Average Pearson’s correlation coefficients of CSF scores with TM-scores. The correlation coefficient of a decoy set is the average coefficient of all targets in that decoy set. In calculating the correlation coefficients, the native structure was excluded. OPUS- CSF has comparable average correlation coefficient with other two potentials. The bold number in each row indicates the best one among the three potential functions for that particular decoy set. For OPUS-CSF, only those results for the combined segment case are listed.

| Decoy sets | OPUS-PSP | GOAP | OPUS-CSF |
| --- | --- | --- | --- |
| 4state_reduced | -0.589 | **-0.694** | -0.667 |
| fisa | -0.282 | -0.347 | **-0.552** |
| fisa_casp3 | -0.095 | -0.221 | **-0.333** |
| hg_structal | -0.752 | **-0.825** | -0.803 |
| ig_structal | -0.779 | -0.865 | **-0.882** |
| ig_structal_hires | -0.832 | -0.885 | **-0.901** |
| I-TASSER | -0.284 | **-0.477** | -0.452 |
| lattice_ssfit | -0.051 | -0.058 | **-0.151** |
| lmds | -0.091 | -0.146 | **-0.342** |
| MOULDER | -0.802 | **-0.886** | -0.863 |
| ROSETTA | -0.343 | **-0.476** | -0.391 |
| Average | -0.521 | -0.632 | -0.624 |

Figure. 1 shows the histogram of standard deviations of the coordinate components of C atoms of the 1^st^ and 5^th^ residues in the CND lookup table for 5-residue segment case. It is clear that the distribution peaks at a very small value indicating that the coordinate components are clustered in a narrow distribution, i.e., the configurational distributions of the 5-residue peptide segments are narrow (Tang and Zhang, 2007), which provides a foundation for the success of OPUS-CSF. The narrow configurational distribution of small peptide fragments is also seen in other studies (Simons et al., 1997). In addition, the average value of the standard deviation is 1.20 Å, in a similar order of magnitude of a single chemical bond length.

**Figure 1.**
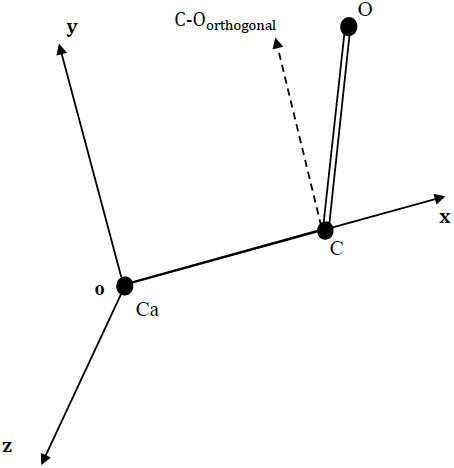
The histogram of standard deviations of the coordinate components in the CND lookup table for 5-residue segment case. The distribution peaks at a very small value of standard deviation indicating that the coordinate components of the 1^st^ and 5^th^ C atoms are clustered in a narrow distribution, i.e., the configurational distributions of the 5-residue peptide segments are narrow. In addition, the average value of the standard deviation is 1.20 Å, in a similar order of magnitude of a single chemical bond length.

It needs to be mentioned that, in the implementation of OPUS-CSF, we assume that the smaller the CSF score, the more likely the structure to be native. This is an approximation because even a native structure may not have a zero CSF score. However, the narrow distributions of standard deviations of the coordinate components of C atoms seem to be in favor of such an approximation. Figure 2 shows the distribution of frequency of sequence repeating in the CND lookup table. Half of the sequences repeat more than 26 times in the distribution. The largest value of X-axis is 29,618 with one sequence.

**Figure 2.**
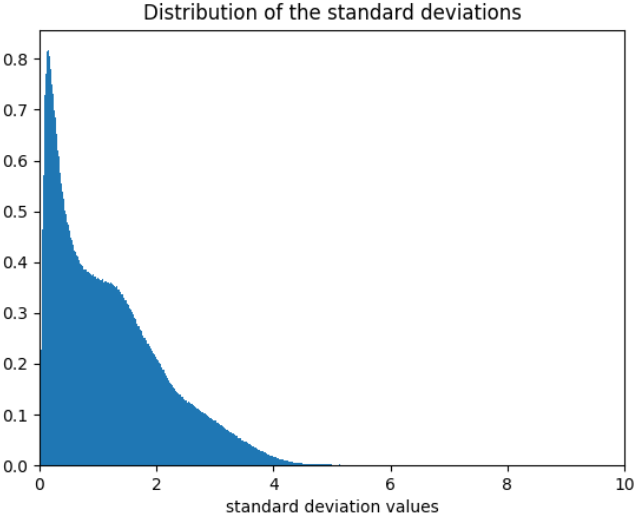
The distribution of frequency of sequence repeating in the CND lookup table. The X-axis is the repeating frequency, and the Y-axis is the number of sequences with particular repeating frequency. Sequences that repeat less than 5 times were omitted in our study. Analysis of this distribution indicates that half of the sequences repeat more than 26 times. The largest value of X-axis is 29,618 with one sequence, but not shown for the purpose of clarity.

We examined OPUS-CSF with different length of residue segment. With the length of segment increases, the ratio of the number of segments that appear more than 5 times to the total number of segments in PDB decreases (Table 3). On the other hand, if the Coverage is defined as the ratio between the number of segments available in CND lookup table and the number of total segments of a test sequence, the average coverage of the 11 decoy sets (totally 278 targets) decreases as the length of segment increases. If a test sequence has less than 20% of its segments available in the CND lookup table, i.e., its coverage is less than 20%, it is regarded as Unknown, then the number of unknowns increases as the length of segment increases. More details of OPUS-CSF on different segment length can be found in Supplemental Information.

**Table 3.** The result of OPUS-CSF built by different length of residue segments. Num_above5 is the number of sequence segments which occur at least five times in PDB. Num_all shows the total number of sequence segments in PDB. The ratio decreases as the length of segments increases.

|  | Num_above5 | Num_all | Num_above5/Num_all |
| --- | --- | --- | --- |
| 5-residues | 1766273 | 2350969 | 0.751 |
| 7-residues | 3736778 | 9544858 | 0.391 |
| 9-residues | 3713506 | 10262243 | 0.362 |
| 11-residues | 3743204 | 10698802 | 0.350 |

Therefore, when working alone, the 5-residue case delivers the best performance in terms of decoy recognition (244 out 278 native recognition in Table 4). However, the Z-scores are better for longer-segment cases. This is probably because the longer segment preserves more sequence homology information.

**Table 4.** The performance of OPUS-CSF based on different length of residue segments on 11 decoys sets. Success numbers are the numbers of native structures that OPUS-CSF correctly recognized from the decoys. Numbers in parentheses (278) is the total number of native structures (or targets) in 11 decoy sets. The Z-scores are the ones calculated based on the CSF scores of the native structures with respect to their decoys. Coverage means the ratio between the number of segments available in CND lookup table and the number of total segments of a target sequence. The table shows the average coverage among 278 targets in 11 decoy sets. Unknowns are the numbers of target sequences that have less than 20% of coverage. For these sequences, OPUS-CSF is not applicable. Note, 5-residue case does not have sequence classified as unknown, while 7-residue case, for example, has 41 out of 278 sequences not applicable for OPUS-CSF. The number of unknown increases slightly as the length of segment increases. Note, in the combined segment case, the longer segments may make no contribution to the CSF score if they are regarded as unknowns. Since5-residue segment case has no unknown, it guarantees OPUS-CSF applicable to all target sequences even in rare ones that all longer segments are regarded as unknown.

|  | 5-residues | 7-residues | 9-residues | 11-residues |
| --- | --- | --- | --- | --- |
| Success numbers | 244 (278) | 218 (278) | 220 (278) | 219 (278) |
| Z-scores | -3.556 | -4.546 | -4.616 | -4.569 |
| Average Coverage | 0.971 | 0.749 | 0.712 | 0.683 |
| Unknowns | 0 | 41 | 45 | 46 |

The construction of CND lookup table ignores the secondary structural elements. We believe this to be an advantage as the prediction of secondary structural elements itself introduces additional uncertainty.

The advantages of OPUS-CSF are obvious. First, the CND lookup table is constructed from the entire PDB, and it contains the information of all allowed configurational information of the native segments (at least for the ones repeated more than 5 times in PDB). Second, the speed of OPUS-CSF is very fast, especially for longer polypeptide chains. This is because the entire chain is scanned once and linearly, it only requires partial mainchain atom coordinates to calculate the CSF score for a structure. Unlike in other potentials such as GOAP (Zhou and Skolnick, 2011) and OPUS-PSP (Lu et al., 2008), no inter-atomic distance needs to be calculated. We want to emphasize that, in modeling protein structures, an empirical potential function or a scoring function, should be fast and accurate. In early stage of modeling, it is advantageous that the scoring function requires minimal amount of structural information. In this regard, OPUS-CSF seems to be a good choice.

## Methods

Scanning through the polypeptide chain with a step size of one residue, we collected small peptide segments with sequence length of 5, 7, 9, and 11 residues and searched for their configurations in the entire PDB. Totally, we downloaded 130,054 PDB structures on June 7, 2017 via ftp://ftp.wwpdb.org/pub/pdb/data/structures/divided/pdb. The sequences that appeared less than 5 times in PDB were discarded.

Here we use 5-residue segment case as an example to illustrate the detail procedure. The ratio of segments that appear more than 5 times to all segments in PDB is 75.1%, which means we can utilize 75.1% information in the whole PDB using 5-residue segments (also see Table 3 in Results and Discussion).

A local molecular coordinate system was defined for every segment using the positions of three main-chain atoms in the middle residue. The origin was set at the Ca atom, the X-axis was defined along the line connecting Ca and C atoms, Y-axis was in the Ca-C-O plane, parallel to component of C-O vector that was perpendicular to the X-axis, and the Z-axis was defined correspondingly (Figure 3).

**Figure 3.**
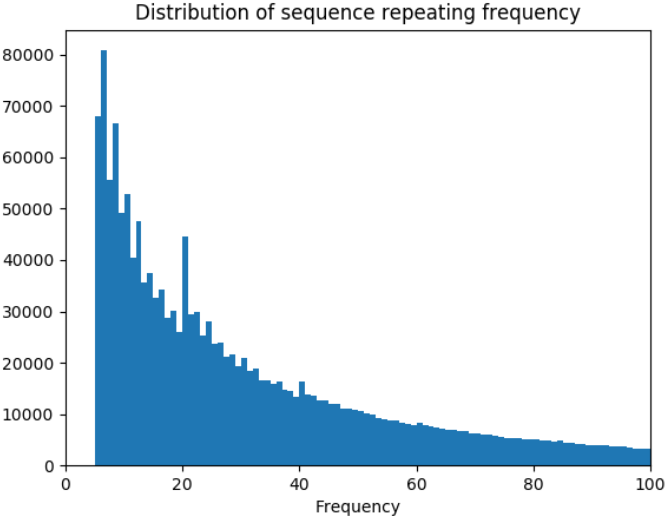
Local molecular coordinate system in OPUS-CSF defined by the mainchain atoms of the 3^rd^ residues. The origin is on Ca atom. The X-axis is along the Ca-C line. Y-axis is in the plan of Ca-C-O atoms, and parallel to the orthogonal projection of C-O vector. Z-axis is defined accordingly.

For a 5-residue segment with a specific sequence, we saved the C atom coordinates of the 1^st^ and 5^th^ residue in the local coordinate system, denoted as (*x*_1_, *y*_1_, *z*_1_) and (*x*_1_, *y*_1_, *z*_1_). And under our assumption, we treated *x*_1_, *y*_1_, *z*_1_, *x*_1_, *y*_1_, *z*_1_ as six independent variables. By scanning through the entire PDB, we generated six independent distributions of these variables, called configurational native distributions (CNDs) of 5-residue segments. We then calculated the means and standard deviations of the distributions and they were kept as the CND lookup table.

For a test structure, we scanned through its sequence with 5-residude-segment. With each segment and its sequence, we looked for the Z-scores of the six independent variables in the CND lookup table. At the end, we added up all the absolute values of Z-scores of all variables for all segments, and it was called CSF score. We assume the polypeptide structure with smallest CSF score has the largest likelihood to be the native structure.

The segments of varying lengths are denoted as 5(1, <u>3</u>, 5), 7(2, <u>4</u>, 6), 9(1, 3, <u>5</u>, 7, 9) and 11(2, 4, <u>6</u>, 8, 10). Here, in segments with the form of 5(1, <u>3</u>, 5), for example, the first number 5 is the segment length, 1,5 in the parenthesis are the residues that we record C atom positional distributions in local coordinate system, <u>3</u> is the residue on which the local coordinate system is defined. For 9(1, 3, <u>5</u>, 7, 9) and 11(2, 4, <u>6</u>, 8, 10), four atoms are used for recording C atom positional distributions, thus totally 12 independent variables are used.

The CSF score can be calculated either based on one particular segment length or by combining all segment length together. In the case of combined segment length, final CSF score is a linear sum of all CSF scores of different segment length. No weighting function is introduced for the contribution of different segment length.

The 11 commonly used decoy sets we used to test OPUS-CSF are the same as those used in GOAP (Zhou and Skolnick, 2011), including decoy sets of 4state_reduced (Park and Levitt, 1996), fisa (Simons et al., 1997), fisa_casp3 (Simons et al., 1997). hg_structal, ig_structal and ig_structal_hires (R. Samudrala, E. Huang, and M. Levitt, unpublished). I-TASSER (Zhang and Zhang, 2010), lattice_ssfit (Samudrala et al., 1999; Xia et al., 2000), lmds (Keasar and Levitt, 2003), MOULDER (John and Sali, 2003) and ROSETTA (Tsai et al., 2003).

## Author Contributions

GX designed the initial concept of OPUS-CSF potential. GX and TM implemented the computational details and thus they made equal contributions to the project. GX and TM also participated in the writing of paper. TZ participated in discussion and contributed to the understanding of the potential function. QW contributed to the understanding of the potential function and participated in the writing of paper. JM supervised the entire project and participated in the writing of paper.

## Acknowledgements

JM thanks support from the National Institutes of Health (R01-GM067801, R01-GM116280), and the Welch Foundation (Q-1512). QW thanks support from the National Institutes of Health (R01-AI067839, R01-GM116280), the Gillson-Longenbaugh Foundation, and The Welch Foundation (Q-1826).

